# LPC 18:2-Driven Apoptosis In Neutrophils Is Non-Inflammatory and Lipid Raft Dependent

**DOI:** 10.64898/2025.12.09.693266

**Authors:** Priyanka Saminathan, Alicia Gibbons, Ian Mathews, Maija Corey, Ashmitaa Logandha, Mahati Rayadurgam, Namratha Nadig, Camille Fang, Neha Reddy, Mousa Vatanmakanian, Sonia Sharma

**Affiliations:** La Jolla Institute for Immunology, La Jolla, CA 92037, United States of America; University of California San Diego, La Jolla, CA, USA; Laboratory for Inflammatory Immune Metabolism, Center for Integrative Medical Sciences, RIKEN, Yokohama, Japan

**Keywords:** Lysophosphatidylcholine (LPC) 18:2, neutrophil apoptosis, reactive oxygen species (ROS), lipid saturation, lipid bilayer integrity, inflammation

## Abstract

Lysophosphatidylcholines (LPCs) are potent bioactive lipids whose fatty acid compositions dictate their immunomodulatory effects. Here, we delineate how unsaturated LPC 18:2 and saturated LPC 16:0 differentially regulate neutrophil survival and inflammatory programs. LPC 18:2 markedly increased reactive oxygen species (ROS) generation and caspase-3/7 activation, mitochondrial membrane depolarization, and cytochrome c release, features consistent with intrinsic apoptosis. In contrast, LPC 16:0 induced robust LDH and HMGB-1 release, indicating membrane rupture and pyroptosis-like death. Bulk RNA sequencing revealed that LPC 16:0 strongly upregulated inflammatory and cytokine gene expression. Disruption of lipid-raft integrity abolished LPC 18:2-induced ROS and apoptosis, underscoring the dependence of these effects on membrane organization. Collectively, these results identify LPC 18:2 as a non-inflammatory, mitochondria-dependent inducer of neutrophil apoptosis, whereas LPC 16:0 promotes inflammatory, lytic death programs. These findings highlight how lipid saturation determines neutrophil fate and immune tone, providing mechanistic insight into how distinct LPC species shape inflammation and tissue injury.

## 1 Introduction

Immune checkpoint blockade (ICB) strategies targeting CTLA-4 and PD-1/PD-L1 molecules have transformed cancer therapy, delivering durable responses across melanoma, non-small cell lung cancer, renal cell carcinoma, and other solid tumors. Unfortunately, ICB treatments frequently cause immune-related adverse events (ICB-irAEs), autoimmune-like toxicities from off-tumor immune activation. ICB-related irAEs span dermatologic, gastrointestinal, endocrine, hepatic, pulmonary, musculoskeletal, neurologic, cardiac, renal, and hematologic toxicities, ranging from mild rashes to life-threatening conditions including myocarditis^1^. ICB-irAEs, particularly severe grade III/IV ICB-irAEs, limit treatment tolerability and are a major factor in patient discontinuation of treatment^2^. Current treatment strategies focus on corticosteroid administration to manage symptoms, which are broadly immunosuppressive and may affect patient treatment response^3^. The precise cellular and molecular drivers of ICB-irAEs remain poorly understood, underscoring a need for research into therapeutics that can serve as target strategies to control pathological immune responses while preserving anti-tumor immunity.

Neutrophils, the most abundant innate immune cells in the blood, are being increasingly appreciated as multifunctional cells with both pro- and antitumoral roles, as well as modulators of ICB efficacy^4,5^ and toxicity^5^. In the tumor microenvironment, neutrophils interact with tumor cells directly, as well as indirectly by the modulation of other immune cells through effector functions including reactive oxygen species (ROS) production, NETosis, and degranulation ^6^. During immunotherapy, high neutrophil-to-lymphocyte ratio (NLR) is associated with worse overall survival and worse progression-free survival across cancer treatment with immune checkpoint blockade agents, as well as with the development of ICB-irAEs. Modulating neutrophil recruitment, function, and fate therefore represents an important strategy to mitigate ICB-irAEs, as well as other conditions with other neutrophil-driven pathology^7,8^. Our medRxiv study is among the first to directly link neutrophilia to severe ICB-irAEs and to association with a specific lipid mediator^9^. We performed longitudinal immune cell profiling and lipidomics across three independent ICB-treated solid tumor patient cohorts and observed that patients with severe ICB-irAEs showed declines in levels of circulating lysophosphatidylcholines LPC 16:0 and LPC 18:2^9^. Low LPC 18:2 levels associated with higher peripheral neutrophil counts in ICB patients and in large healthy population cohorts, implicating an LPC 18:2–neutrophil homeostasis axis in ICB-irAE risk.

LPCs comprise a class of circulating bioactive lipid mediators distinguished by their acyl chain compositions, and dysregulation of their circulating concentrations has been observed in a myriad of different diseases^10–16^. LPCs directly reprogram immune cells, including neutrophils and macrophages, as well as endothelial cells, *in vitro*, with phenotypes specific to the acyl chain composition of the LPC species studied^16,17^. *In vivo*, our group observed that intraperitoneal delivery of LPC 18:2 alleviated colonic neutrophilia and prevented neutrophil-mediated tissue damage in both ICB-irAE colitis and dextran sodium sulfate (DSS) colitis models^18^. We also observed that upon colitis induction by either ICB-irAEs or DSS, circulating LPC 18:2 levels dropped^9^. Together, these data support a causal, protective role for LPC 18:2 across multiple inflammatory models relevant to ICB toxicity^9^. These findings warrant dissecting neutrophil fate in ICB and advancing lipid-immune strategies to curb ICB-irAEs without compromising antitumor immunity.

While we observed that both LPC 16:0 and LPC 18:2 were linked to lower ICB-irAE risk, only LPC 18:2 was inversely associated with patient blood neutrophilia^9^. Unlike LPC 18:2, supplementation with LPC 16:0 yielded no protective phenotype in murine colitis models^9^. We hypothesize that this divergence reflects acyl-chain specific control of neutrophil fate. Specifically, we posit that LPC 18:2 signals through lipid-raft architecture to trigger a transcriptionally silent, intrinsic apoptosis program in primary human neutrophils, whereas LPC 16:0 promotes pro-inflammatory gene transcription. We test this by integrating bulk RNA-seq with functional readouts of apoptosis and mitochondrial integrity in human neutrophils, alongside perturbation of lipid raft order. These data explain our observed clinical pattern in ICB-irAE patients and support a targeted LPC 18:2 strategy to limit neutrophil-mediated ICB-irAEs.

## 2 Methods

### 2.1 Isolation of human neutrophils

The Miltenyi Whole Blood Neutrophil Isolation Kit (Miltenyi Biotec, Cat# 130-104-434) was used to negatively isolate human neutrophils from whole blood from healthy human donors according to the manufacturer’s instructions. Briefly, whole blood was incubated with reconstituted antibody-labeled beads at room temperature for 5 minutes then placed in a magnetic separator for 30 minutes. Following incubation, the supernatant containing neutrophils was collected, and red blood cell lysis was performed for 5 minutes at room temperature. Neutrophils were subsequently resuspended in RPMI 1640 Medium (ATCC modification) (ThermoFisher Scientific Cat# A1049101) supplemented with 5% heat-inactivated FBS and maintained at 37 °C until use.

### 2.2 ROS assay

A total of 0.4 × 10□ cells were seeded per well in a V-bottom 96-well tissue culture-treated plate. Cells were washed twice with FBS-free RPMI 1640 Medium (ATCC modification), then incubated with 100 µL of 10 µM dihydrorhodamine 123 (DHR, ThermoFisher Scientific Cat# D632) for 20 minutes at 37°C to assess reactive oxygen species (ROS) production. After incubation, 100 µL of the respective test condition compounds (prepared at 2× concentration) were added directly to each well, resulting in a final volume of 200 µL. Cells were incubated under these conditions for 45 minutes at 37°C. After treatment, plates were placed on ice for 5 minutes to halt cellular processes. Cells were then centrifuged at 300 × g for 5 minutes, and supernatants were discarded. In order to stain for apoptotic neutrophils, samples were washed twice with the Annexin V binding buffer (BioLegend) and stained using the Pacific Blue labeled Annexin V (BioLegend) as per instructions provided by the manufacturer (15 minutes at room temperature in the dark). Samples were finally resuspended in 200 µL of FACS buffer (1×PBS with 2% FBS + 0.05% Sodium Azide) containing propidium iodie (viability (PI, 1:200 dilution). Samples were immediately analyzed on an BD LSRFortessa™ X-20 Cell Analyzer (BD Biosciences) using the BD FACSDiva™ Software.

### 2.3 Caspase 3/7 activity assay

A total of 0.4 × 10□ cells were seeded per well in a V-bottom 96-well tissue culture-treated plate. Cells were washed twice with FBS-free RPMI 1640 Medium (ATCC modification). Following washes, cells were treated with 200 µL of the respective experimental condition compounds for the designated time periods. At the end of incubation, plates were placed on ice for 5 minutes to halt cellular activity. Cells were subsequently centrifuged at 300 × g for 5 minutes. Cell pellets were then resuspended in 100 µL of a 1:1000 dilution of Caspase-3/7 detection reagent (as per the manufacturer’s instructions (ThermoFisher Scientific Cat# C10740) prepared in a pre-warmed (37°C) FACS buffer (1×PBS with 2% FBS). Cells were incubated at 37°C for 25 minutes. Following incubation, 100 µL of SYTOX solution (1:500 dilution in FACS buffer) was added to each well and incubation was continued for an additional 5 minutes. Samples were analyzed using a BD® LSR II Flow Cytometer (BD Biosciences) using the BD FACSDiva™ Software.

### 2.4 LDH assay

Cytotoxicity was measured using the CyQUANT™ LDH Cytotoxicity Assay Kit (ThermoFisher Scientific, Cat# C20300) according to the manufacturer’s instructions. Cells were plated in 96-well V-bottom plates and incubated at 37 °C for 30 minutes. After treatment, 50 µL of culture supernat nt from each well was transferred to a clear flat-bottom 96-well plate. Cells were lysed with 10× Lysis Buffer. Reaction Mixture 50 µL was added to each well and incubated at room temperature for 30 minutes, protected from light. The reaction was stopped by adding 50 µL of Stop Solution, and absorbance was measured at 490 nm with 680 nm as reference.

### 2.5 Water-soluble tetrazolium salt-1 (WST-1) assay

Cells were seeded in 96-well plates at a density of 0.4 × 10□ cells in 100 µL of FBS-free RPMI 1640 Medium (ATCC modification), with or without test compounds, and incubated at 37 °C for 2 hours. Following incubation, 10 µL of WST-1 reagent (Cayman Chemical Company, Cat# 10008883) was added to each well, and plates were gently mixed on an orbital shaker for 1 minute. Cells were then incubated for 4 hours at 37 °C. Prior to measurement, plates were mixed again for 1 minute. Absorbance was measured at 450 nm using a microplate reader.

### 2.6 Bulk RNA sequencing

Cells were seeded in deep-well 96-well plates at a density of 2 × 10□ cell in 1 mL of FBS-free RPMI 1640 Medium (ATCC modification), with or without test compounds (LPC 16:0 or LPC 18:2 at 10 μM), and incubated at 37 °C for 2 hours. Total RNA was extracted from 1.5 million cells using the Direct-zol RNA Miniprep Kit (Zymo Research, Cat# R2050). Libraries were prepared using the NEB Ultra II Directional RNA Library Prep Kit for Illumina with the NEBNext Poly(A) mRNA Magnetic Isolation Module. 20 ng total RNA was used for each sample and libraries underwent 16 cycles of PCR amplification. Agilent Universal Human Reference RNA was prepared as a process control. Libraries were sequenced on the Illumina Novaseq 6000 using a 50×50 run configuration at a targeted depth of 25M (50M paired-end) reads.

### 2.7 RNA-seq analysis pipeline

The paired-end reads that passed Illumina filters were filtered for reads aligning to tRNA, rRNA, adapter sequences, and spike-in controls. The reads were then aligned to GRCh38 reference genome and Gencode v27 annotations using STAR (v 2.6.1)^19^. DUST scores were calculated with PRINSEQ Lite (v0.20.3)^20^ and low-complexity reads (DUST >4) were removed from the BAM files. The alignment results were parsed via the SAMtools^21^ to generate SAM files. Read counts to each genomic feature were obtained with featureCount (v 1.6.5)^22^. After removing absent features (zero counts in all samples), the raw counts were then imported to R/Bioconductor package DESeq2 (v 1.24.0)^23^ to identify differentially expressed genes among samples. P-values for differential expression were calculated using the Wald test for differences between the base means of two conditions. These P-values were then adjusted for multiple test correction using the Benjamini Hochberg algorithm^24^. Gene set enrichment analysis was run using the ‘GseaPreranked’ method with ‘classic’ scoring scheme. MSigDB Gene sets were downloaded from the MSigDB website. Rank files for each DE comparison of interest were generated by assigning a rank of negative log10(pValue) to genes with log2FoldChange greater than zero and a rank of positive log10(pValue) to genes with log2FoldChange less than zero.

### 2.8 Lipid-raft disruption

Cells were seeded at 0.4 × 10□ cells per well in a 96-well V-bottom plate, centrifuged and then resuspended with MβCD at final concentrations of 500 µM or 1 mM to disrupt cholesterol-rich lipid raft microdomains. A total of 100 µL of the MβCD solution was added per well, and cells were incubated at 37 °C for 35 minutes prior to treatment with test compounds.

### 2.9 Cytochrome c release assay

To test for cytochrome c release, a widely used readout of mitochondrial integrity and apoptotic cell death, post-treatment with test compounds, cells were fixed with 100 µl/well 4% PFA for 10 minutes at room temperature. Following fixation, cells were washed with 1×PBS, and then permeabilized with 0.1% PBS-TritonX-100 for 15 minutes at room temperature. After incubation, cells were washed with 1×PBS and blocked with 10% Goat serum in 1×PBS for 30 minutes at room temperature. Next, the cells were stained with Cytochrome C Monoclonal Antibody FITC (eBioscience, Cat# 11-6601-82) at a 1:100 concentration in the blocking buffer and left to incubate for 8 hours at 4 °C. Cells were then washed and resuspended in 1×PBS and analyzed using the BD® LSR II Flow Cytometer (BD Biosciences)

### 2.10 Mitochondrial potential assay (MitoTracker Red assay)

The MitoTracker Red assay is used to assess mitochondrial membrane potential, a key indicator of mitochondrial health. The dye accumulates in active mitochondria in a potential-dependent manner, so loss of fluorescence reflects mitochondrial depolarization and subsequent dysfunction, which occurs as a result of intrinsic apoptosis. Briefly, after incubation with test compounds, neutrophils were spun down, supernatant removed, and the cells were resuspended with 100 µl of RPMI 1640 Medium (ATCC modification) containing a final concentration of 50nM MitoTracker Red CMXRos (Thermo Scientific, Cat# M7512). Cells were then incubated at 37°C for 35 minutes. Finally, cells were resuspended in FACS buffer (1×PBS + 2% FBS) containing 1:8000 DAPI and then analyzed on the BD LSRFortessa™ Cell Analyzer.

### 2.11 Statistical analysis

Statistical analyses for large datasets (bulk RNA-seq data) were performed in R. Statistical tests for *in vitro* studies were performed in GraphPad Prism. Experiments were independently replicated at a minimum of 3 times.

## 3 Results

### 3.1 LPC 16:0 induces pyroptosis unlike LPC 18:2

Plasma lipidomic profiling from ICB-treated patients revealed that individuals without ICB-irAEs displayed significantly higher levels of circulating LPC species compared to those who did develop ICB-irAEs, suggesting a protective association between LPC abundance and systemic inflammation^9^. Previously, we have shown that LPC 18:2 supplementation abrogates ICB-irAE development in mouse models by limiting neutrophilia under ICB-induced inflammatory conditions^9^. Among individual species, LPC 18:2 correlated inversely with neutrophil counts in our ICB-patient cohorts, while LPC 16:0 showed no correlation with neutrophil counts. This observation instigated our study of the distinct effects of saturated and unsaturated LPC species on neutrophil fate *in vitro*. To investigate the effects of saturated and unsaturated LPC species on neutrophils, primary neutrophils were isolated from the peripheral blood of healthy human patients and exposed to physiologically relevant concentrations of LPC species (10-30 µM)^17^. We focused on LPC 16:0 and LPC 18:2, the two most abundant LPC species in the plasma, both of which were inversely associated with severe ICB-irAEs in our patient cohort, and the latter of which we also observed to be inversely associated with ICB-irAE-associated patient neutrophilia^9,17^. Cellular metabolic activity was quantified using the WST-1 assay (Fig. 1a), which measures extracellular reduction of tetrazolium salts by plasma-membrane NADPH oxidoreductases and intact mitochondrial redox systems. LPC 16:0 10 µM treatment resulted in a significant increase (approx. 25% increase) of WST-1 expression. Given that WST-1 depends on mitochondrial and cytosolic dehydrogenase enzyme activity and considering the terminally differentiated (non-dividing) nature of neutrophils, this increase in WST-1 signal may be indicative of increased metabolic activity^25,26^. LPC 16:0 30µM treatment significantly decreased WST-1 reduction by approx. 90.3%, consistent with rapid loss of redox capacity and disruption of membrane integrity^27^. This significant decrease in WST-1 expression suggests that LPC 16:0 triggers an early metabolic collapse consistent with lytic cell death. In contrast, LPC 18:2 treatment produced only a modest, non-significant increase in WST-1 expression, observed only at 30□µM, while treatment at 10□µM caused no change in WST-1 levels. Treatment with either LPC species increased caspase 3/7 activity, suggesting initiation of apoptosis (Fig. 1b). LPC 16:0 (10 µM) induced significant apoptosis in neutrophils, while LPC 16:0 (30 µM) treatment induced loss of viability through the induction of necrosis (Fig. 1c, gating scheme Supplemental Fig. 1). Caspase 3/7 activity was significantly elevated in LPC 18:2 30 µM treated neutrophils (Fig. 1b).We next examined additional markers of cell death and stress, specifically lactate dehydrogenase (LDH) and high-mobility group box 1 (HMGB1), which are also indicators of pyroptosis. Consistently, LDH release at 1.5 hours post treatment increased nearly 10-fold in LPC 16:0-treated neutrophils, compared with only a <2-fold increase following LPC 18:2 treatment, confirming extensive membrane rupture (Fig. 1d). At 2 hours post-treatment, high mobility group box-1 (HMGB1) release, a hallmark of pyroptotic or necrotic cell lysis, was elevated 1.84-fold following LPC 16:0 (10 µM) and 1.66-fold following LPC 16:0 (30 µM) treatment in comparison to unstimulated cells (Fig. 1e). In contrast, LPC 18:2 induced no detectable HMGB1 expression at 10 µM and only an approximately 1.2-fold increase at 30 µM. To further investigate the cellular stress responses underlying the observed changes in WST-1, LDH, HMGB1 release, and apoptosis, we measured reactive oxygen species (ROS) generation in neutrophils. ROS levels were quantified in live cells by flow cytometry, with late-apoptotic cells (Annexin V+ neutrophils) gated out, allowing assessment of oxidative stress specifically in live neutrophils (Fig. 1f, gating scheme Supplemental Fig 1). Consistently with a previously published study^17^, we observed that LPC 18:2 significantly increased ROS production (12.7 fold increase) in contrast to the modest (and statistically insignificant) increase (1.67 fold at 10 μM and 2.07 fold at 30 μM) with LPC 16:0 (Fig. 1f). Together, these data demonstrate that LPC 16:0 promotes rapid, lytic neutrophil death with little ROS production, while LPC 18:2 favors a modest caspase-3/7 activation, robust ROS production and preserved viability.

**Figure 1.**
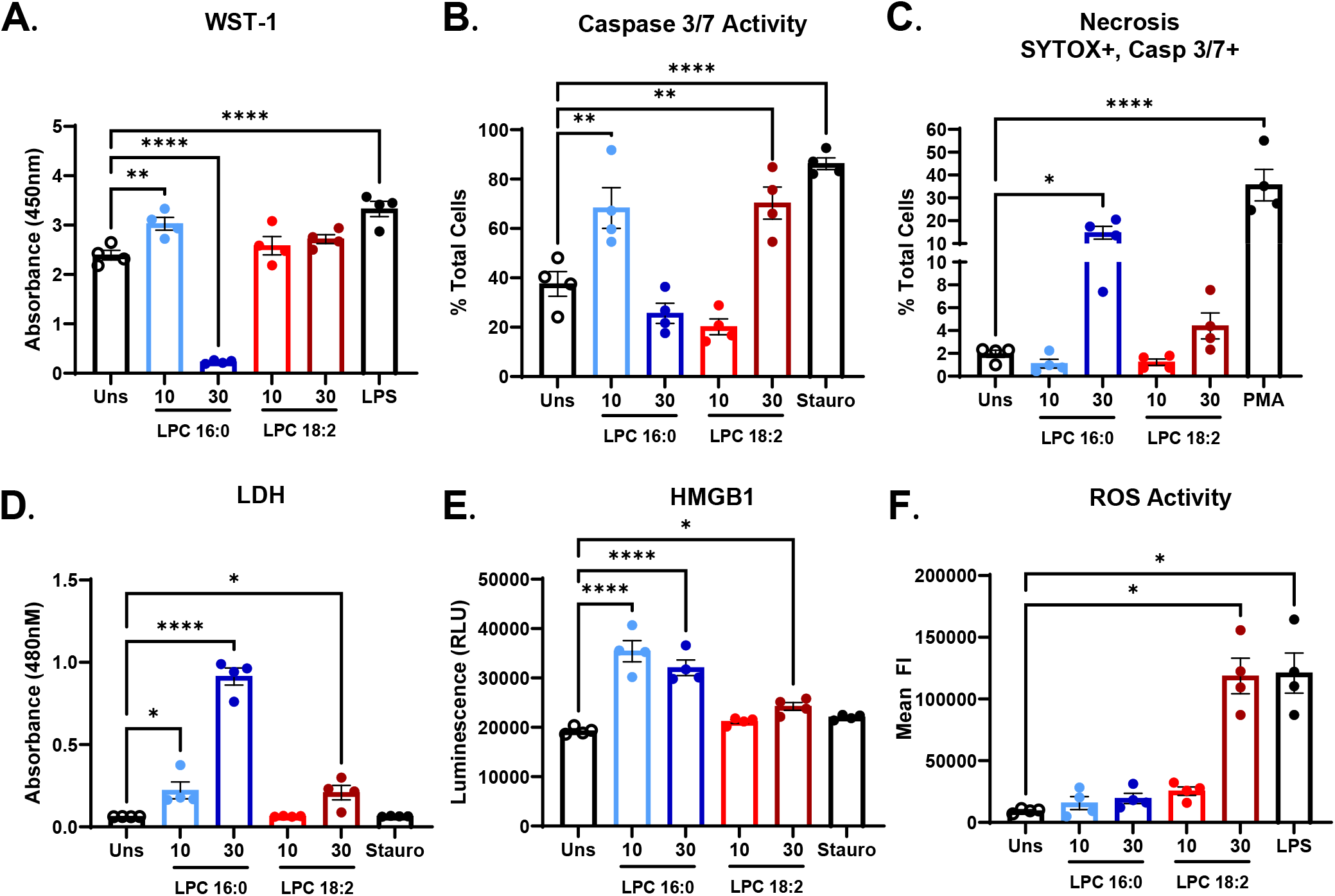
LPC 16:0 and LPC 18:2 differentially modulate human neutrophils. (A) Human neutrophil cellular metabolic activity measured by WST-1 assay after treatment with LPC 16:0, LPC 18:2, or Lipopolysaccharide (LPS). (B) Human neutrophil apoptosis (cleaved caspase-3/7) after treatment with LPC 16:0, LPC 18:2, or Staurosporine. (C) Human neutrophil necrosis (Sytox+ caspase-3/7 dual stain) after treatment with LPC 16:0, LPC 18:2, or Phorbol 12-myristate 13-acetate (PMA). (D) Lactate Dehydrogenase (LDH) release by human neutrophils after treatment with LPC 16:0, LPC 18:2, or Staurosporine. (E) High-Mobility Group Box 1 (HGMB1) release by human neutrophils after treatment with LPC 16:0, LPC 18:2, or Staurosporine. (F) ROS induction in human neutrophils treated with LPC 16:0, LPC 18:2, or LPS. Statistics: Ordinary one-way ANOVA with Dunnett’s test (A-F); in all panels, bars = mean, error = SEM. (*p ≤ 0.05, **p ≤ 0.01, ***p ≤ 0.001, ****p ≤ 0.0001).

### 3.2 Bulk-sequencing reveals LPC 16:0 more pro-inflammatory than LPC 18:2

To investigate the molecular basis of the differential neutrophil responses to LPC 16:0 and LPC 18:2, we performed bulk RNA sequencing on isolated human donor neutrophils (n = 3) following 45 minute treatment with each LPC species (at 10 µM). Principal component analysis (PCA) revealed clear separation between LPC 16:0-treated cells and unstimulated controls, whereas LPC 18:2-treated cells clustered closely with controls (Fig. 2a). This pattern indicates transcriptional reprogramming in response to LPC 16:0 treatment, but minimal gene expression changes with LPC 18:2 treatment, suggesting LPC 18:2 acts in a post-transcriptional manner at this concentration. Differential expression analysis identified 81 upregulated and 19 downregulated genes in LPC 16:0-treated cel s relative to control (FDR < 0.01, |log□FC| > 1) (Fig. 2b). No gene transcripts were observed to be significantly differentially expressed after LPC 18:2 treatment (Fig. 2c). Top LPC 16:0-upregulated genes included inflammatory mediators *FOS, FOSB, PTGS2, CXCL8, CCL3L3*, and *DUSP1*, which together regulate neutrophil activation, cytokine production, and survival (Fig. 2d). Top LPC 16:0-downregulated genes included *DEFA3, CD300E*, and *MPO*, with functions in degranulation and oxidative burst (Fig. 2e). Other top downregulated genes including *HBA2, HBB*, and *IL2RB* likely represent erythroid or lymphocyte cell or RNA contamination during neutrophil isolation (Fig. 2e). WikiPathways, Reactome, PID, and KEGG pathway analyses highlighted enrichment of inflammatory and stress-response pathways in LPC 16:0-treated cells (Fig. 2f). Upregulated programs included TNF signaling, IL-1 family signaling, cytokine-cytokine receptor interaction, MAPK activation, and NF-κB-dependent responses, confirming a broad transcriptional inflammatory signature (Fig. 2f). Direct comparison between LPC 16:0 and LPC 18:2 conditions highlighted opposing regulation of inflammatory and cell viability programs, mirroring the functional assays (Fig. 1) where LPC 16:0 induced higher lytic death (LDH expression) and HMGB1 release in when compared with LPC 18:2.

**Figure 2.**
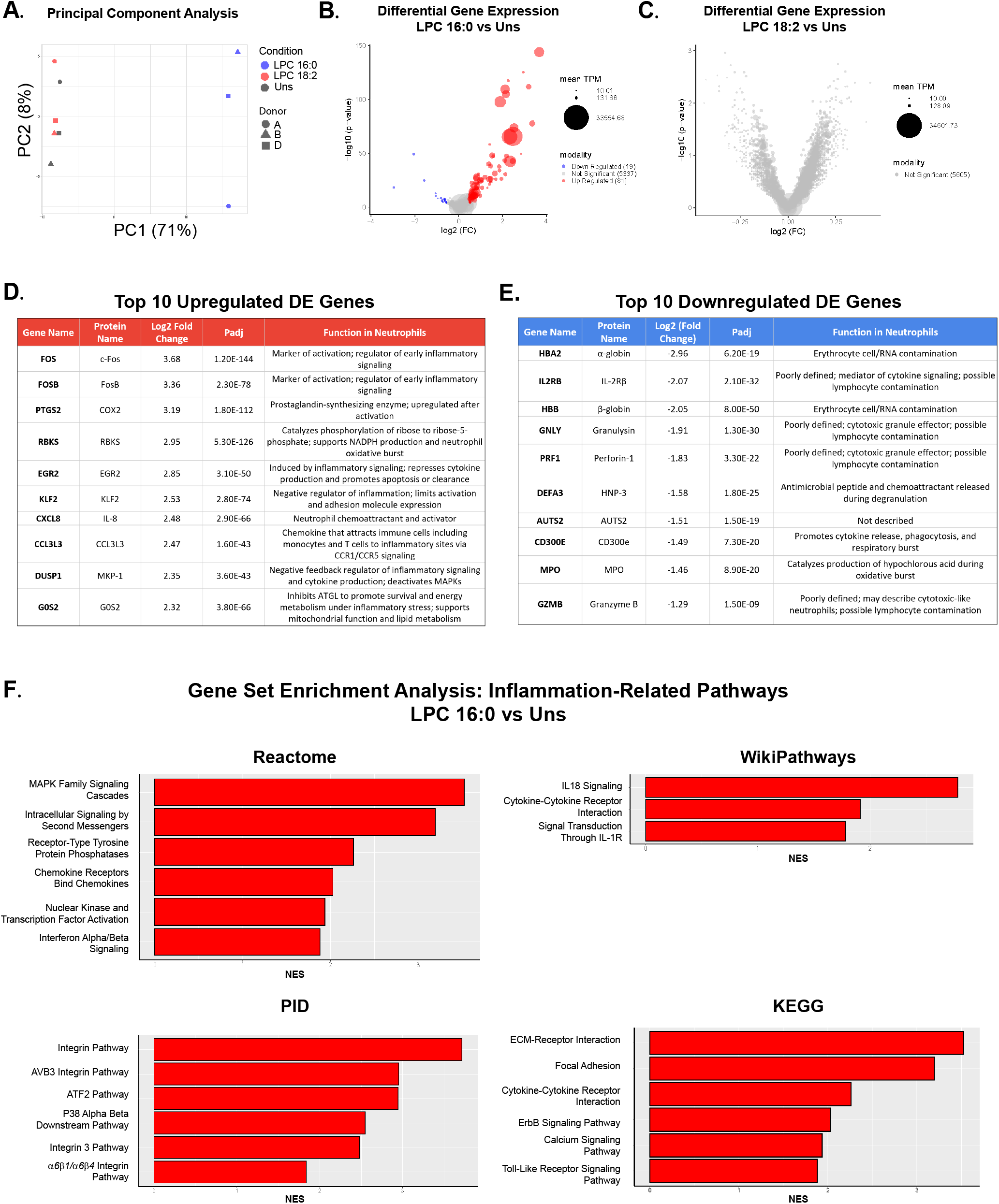
LPC 16:0 but not LPC 18:2 upregulates inflammatory neutrophil gene expression. (A) Principal component analysis (PCA) of human neutrophils treated with media vehicle (unstimulated), 10 µM LPC 16:0, or 10 µM LPC 18:2. (B) Volcano plot of differentially expressed genes (DEGs) between LPC 16:0-treated neutrophils and unstimulated cells (DESeq2). DEG threshold: FDR < 0.01, |log□FC| > 1. (C) Volcano plot of DEGs between LPC 18:2-treated neutrophils and unstimulated cells (DESeq2). DEG threshold: FDR < 0.01, |log□FC| > 1. (D) Top 10 significantly upregulated genes after LPC 16:0 treatment. (E) Top 10 significantly downregulated genes after LPC 16:0 treatment. (F) Inflammation-related gene pathways upregulated after LPC 16:0 treatment identified by gene set enrichment analysis (Reactome, Wikipathways, PID, and KEGG).

### 3.3 LPC 18:2 induces intrinsic apoptosis

LPC 18:2 can serve as a substrate for the secreted enzyme Autotaxin (ATX, also known as lysophospholipase□D), which catalyses its conversion into lysophosphatidic acid (LPA□18:2). LPA is a bioactive lipid mediator implicated in inflammation and neutrophil activation^28^. Although LPC□18:2 can be enzymatically converted to the bioactive lipid LPA□18:2, we observed that neutrophil apoptosis was driven specifically by LPC 18:2 and not LPA 18:2 as seen through the induction of caspase 3/7 activity in our experimental system (Fig. 3a). Apoptosis is a regulated form of programmed non-inflammatory cell death characterized by caspase activation, mitochondrial outer membrane permeabilization, and nuclear condensation, all occurring without plasma membrane rupture^29^. Two major apoptotic pathways exist: the extrinsic pathway, triggered by signaling through plasma membrane death receptors, and the intrinsic (mitochondrial) pathway, which is governed by mitochondrial integrity and cytochrome c release into the cytosol^29^. To determine whether LPC 18:2 treated neutrophils undergo intrinsic apoptosis, we measured cytochrome c release (Fig. 3b) and mitochondrial membrane potential (Fig. 3c and d) following exposure to 30µM LPC 18:2. LPC 18:2 markedly increased cytosolic cytochrome c levels (Fig. 3b, gating scheme Supplemental Fig 1). In a separate experiment, we assessed neutrophil mitochondrial membrane potential upon LPC 18:2 treatment. By gating out dead cells (DAPI□) via flow cytometry, we specifically assessed changes in mitochondrial membrane potential in live neutrophils following LPC treatment. Mitochondrial potential was quantified using MitoTracker fluorescence intensity. During neutrophil apoptosis, mitochondrial potential typically decreases, as observed with staurosporine treatment (Fig. 3c and 3d, *left*, gating scheme Supplemental Fig 1). Upon LPC 18:2 30 µM treatment, three distinct MitoTracker fluorescence peaks emerged, which we categorized as High (Hi), Medium (Mid), and Low (Lo) mitochondrial potential, representing three tiers of mitochondrial polarization (Fig. 3c). Quantification of the DAPI□ population revealed that LPC□18:2 30 µM significantly decreased mitochondrial potential, evidenced by a significant reduction in the “Hi” population and a corresponding increase in the “Mid” population. Interestingly, the mean fluorescence intensity (MFI) of MitoTracker within the remaining Hi population increased with LPC 18:2 30 µM treatment which was not observed with staurosporine treatment (Fig. 3d, *right*). This is potentially indicative of elevated ROS activity in the live, non-apoptotic neutrophil subset, as observed in Fig. 1f. With increases in caspase 3/7 activity, an executioner caspase downstream of cytochrome c signaling, these findings indicate that LPC 18:2 triggers mitochondria-dependent intrinsic apoptosis. This response contrasts sharply with LPC 16:0, which promoted early metabolic collapse and lytic cell death (Fig. 1). The ability of LPC 18:2 to engage intrinsic apoptotic machinery while preserving membrane integrity suggests a protective, resolution-oriented death program in neutrophils, aligning with its non-inflammatory profile observed in ICB patient plasma.

**Figure 3.**
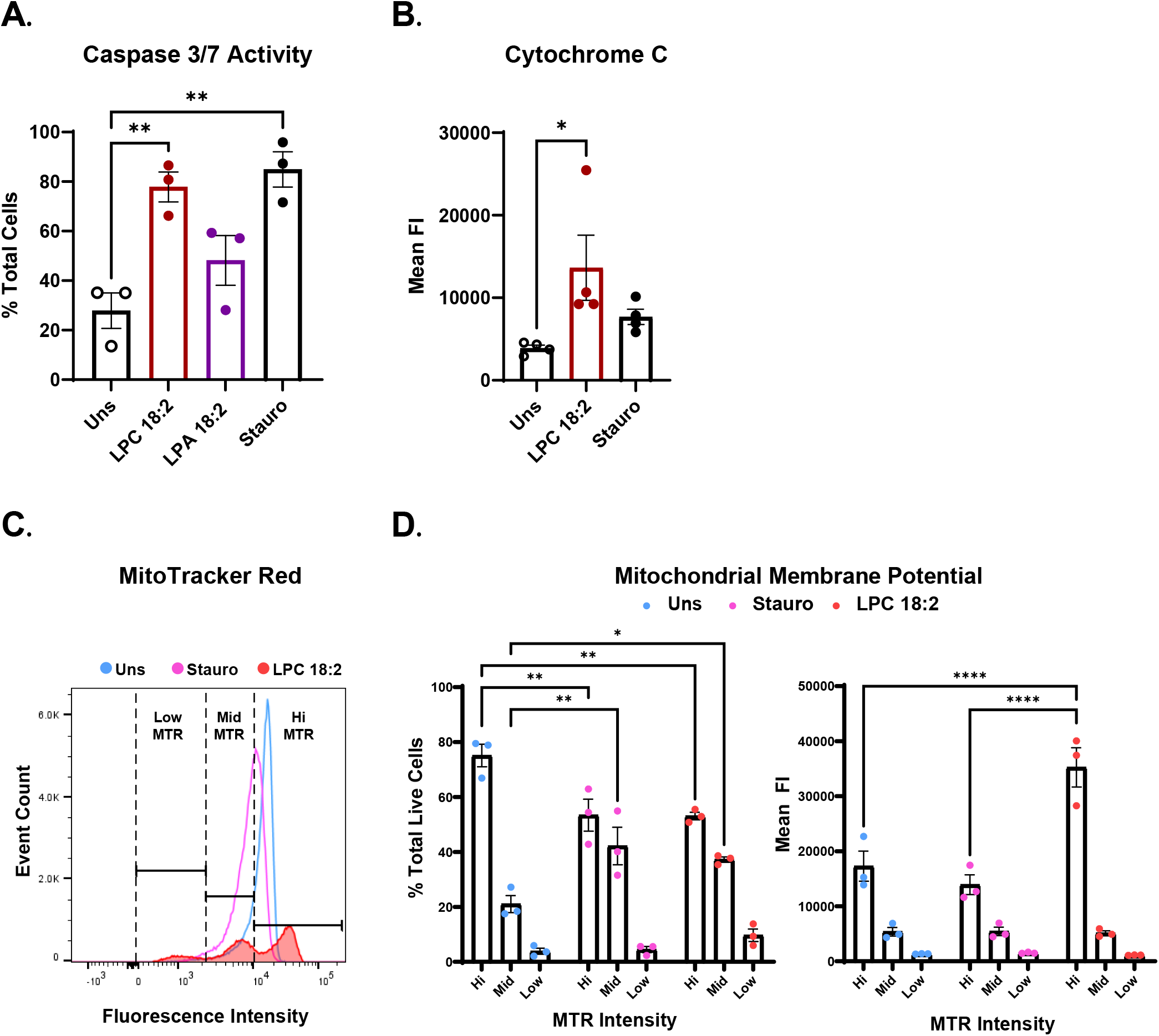
LPC 18:2 induces intrinsic apoptosis. (A) Human neutrophil apoptosis (cleaved caspase-3/7) after treatment with LPC 18:2, LPA 18:2, or Staurosporine. (B) Cytochrome c release by human neutrophils after treatment with LPC 18:2 or Staurosporine. (C) MitoTracker Red fluorescence intensity of human neutrophils after treatment with LPC 18:2 or Staurosporine. Three populations were observed after LPC 18:2 treatment (low MTR, mid MTR, and hi MTR). (D) Quantification of % total live cells and mean FI of populations from 3c. Statistics: Ordinary one-way ANOVA with Dunnett’s test. (A, B), two-way ANOVA with Tukey’s test (D); in all panels, bars = mean, error = SEM. (*p ≤ 0.05, **p ≤ 0.01, ***p ≤ 0.001, ****p ≤ 0.0001).

### 3.4 Lipid raft stability is integral for LPC 18:2 driven ROS activity and apoptosis

Lipid rafts are cholesterol- and sphingolipid-enriched membrane microdomains that serve as organizing platforms for immune receptor signaling and lipid–protein interactions^30^. LPC 16:0 has been shown to incorporate into and signal through lipid rafts to regulate calcium flux and NADPH oxidase activity in leukocytes^31,32^. To test whether LPC 18:2 signaling similarly depends on raft integrity, neutrophils were pretreated with methyl-β-cyclodextrin (MβCD) to deplete membrane cholesterol and thereby disrupt lipid raft order^33^. Raft disruption completely abolished LPC 18:2-induced ROS generation and viability loss measured by DAPI staining (Fig. 4a, b). MβCD pretreatment likewise eliminated LPC 18:2-mediated caspase-3/7 activation (Fig. 4c) and cell necrosis defined by Sytox+, Caspase-3/7+ staining (Fig. 4d). These findings demonstrate that both oxidative and apoptotic responses to LPC 18:2 treatment require intact lipid-raft organization. Together, these results identify lipid rafts as essential scaffolds for LPC 18:2-neutrophil engagement, suggesting that the polyunsaturated LPC species signal through lipid raft-dependent surface or membrane proximal pathways rather than by passive diffusion or interaction with a non-lipid raft-localized protein receptor.

**Figure 4.**
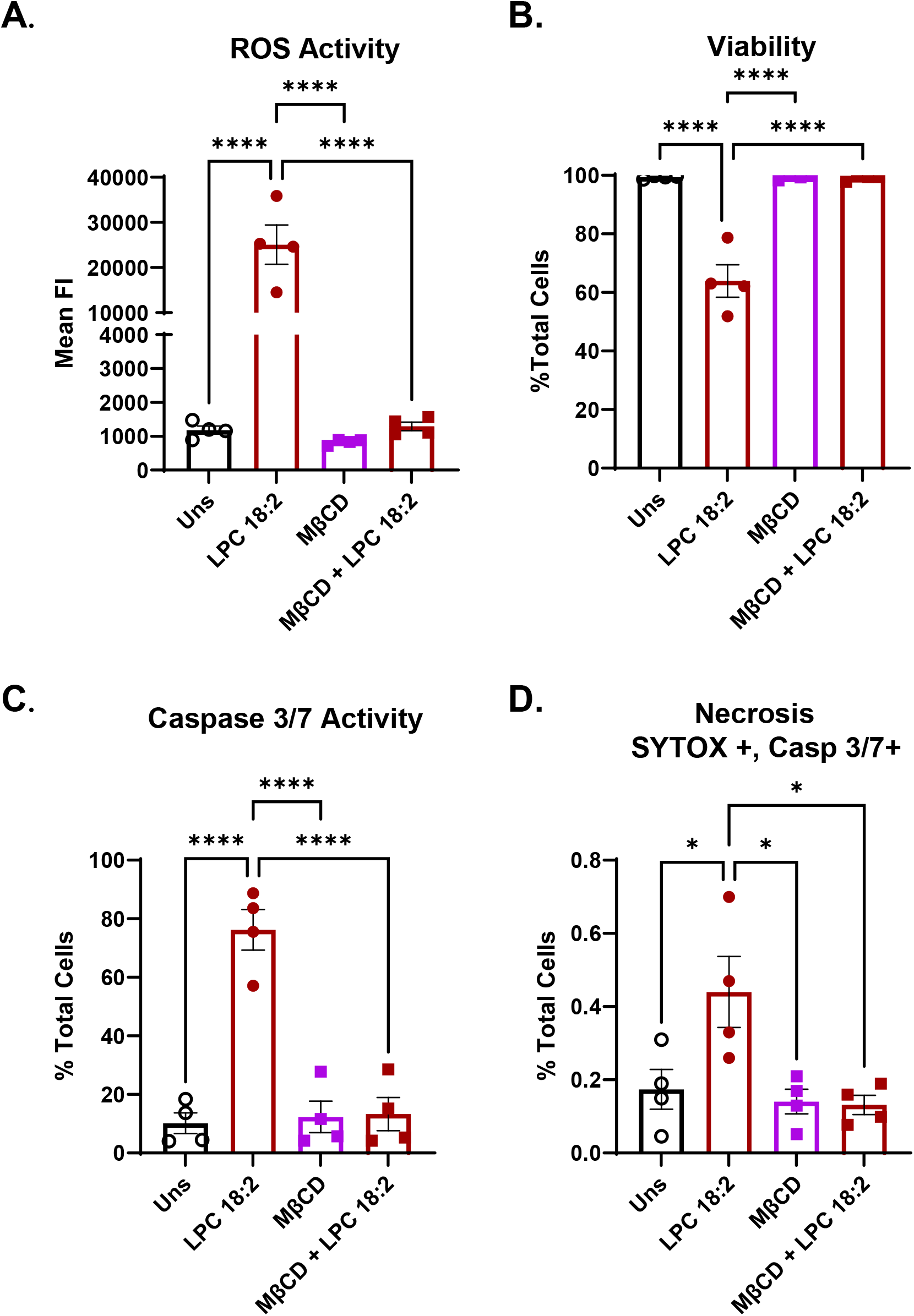
LPC 18:2-induced ROS and apoptosis is dependent on cholesterol and lipid rafts. (A) ROS induction in human neutrophils treated with LPC 18:2, Methyl-β-cyclodextrin (MβCD), or LPC 18:2 and MβCD. (B) Viability (DAPI-%) of human neutrophils after treatment with LPC 18:2, Methyl-β-cyclodextrin (MβCD), or LPC 18:2 and MβCD. (C) Human neutrophil apoptosis (cleaved caspase-3/7) after treatment with LPC 18:2, Methyl-β-cyclodextrin (MβCD), or LPC 18:2 and MβCD. (D) Human neutrophil necrosis (Sytox+ caspase-3/7 dual stain) after treatment with LPC 18:2, Methyl-β-cyclodextrin (MβCD), or LPC 18:2 and MβCD. Statistics: Ordinary one-way ANOVA with Tukey’s test (A-D); in all panels, bars = mean, error = SEM. (*p ≤ 0.05, **p ≤ 0.01, ***p ≤ 0.001, ****p ≤ 0.0001).

## 4 Discussion

In this study, we identify LPC 18:2 as a lipid mediator that shapes neutrophil fate and function in a distinct manner compared to LPC 16:0. Treatment of primary human neutrophils with physiologically relevant concentrations of LPC 18:2 triggered transcriptionally silent ROS production and intrinsic apoptosis via a cholesterol- and lipid raft–dependent mechanism. These findings suggest that LPC 18:2 may help constrain neutrophil-mediated inflammation and tissue damage by inducing intrinsic apoptosis, thus providing the beginnings of a mechanistic framework for understanding how LPC 18:2 may protect against severe ICB-irAEs through neutrophil reprogramming.

A notable observation of this work is the acyl-chain-specific effects of LPC species on neutrophil function. LPC species are established to have acyl chain-dependent effects: saturated LPC species are generally inflammatory, while polyunsaturated species are generally non-inflammatory or inflammation-resolving^34^. Work by Ojala et al. showed that in primary neutrophils, unsaturated LPC species elicit a stronger respiratory burst than saturated species. Unsaturated species instead cause greater calcium influx and disruption of membrane permeability^17^. Here, we corroborate that LPC 16:0, but not LPC 18:2 treatment, disrupts neutrophil membrane permeability. Specifically, we observed that LPC 18:2 induced higher ROS but lower levels of apoptosis than LPC 16:0 at the same treatment concentrations. LPC 18:2 treatment further induced mitochondrial membrane potential loss and cytochrome c release, indicative of activation of the intrinsic apoptosis pathway. Furthermore, LPC 16:0 treatment, but not LPC 18:2 treatment, robustly activated proinflammatory pathways, as evidenced by transcriptional upregulation of immediate-early genes (*FOS, JUN*), and inflammatory cytokines including *CXCL8*. Our observation of *CXCL8* induction by LPC 16:0 treatment is consistent with a previous report in human endothelial cells showing that LPC 16:0 treatment, but not LPC 18:2 treatment, promoted *CXCL8* (IL-8) gene expression^35^. In addition, our observation that LPC 16:0 treatment increased *PTGS2* (COX2) gene expression corroborates a study demonstrating that LPC 16:0 treatment, but not LPC 18:2 treatment, upregulated *PTGS2* expression in endothelial cells^36^. In the referenced study, treatment with both compounds increased COX2 protein expression, providing additional evidence that LPC 18:2 acts post-transcriptionally^36^. Altogether, our data demonstrate that LPC 16:0 treatment promoted a proinflammatory transcriptional program, whereas LPC 18:2 treatment resulted in transcriptionally quiet apoptosis, reinforcing the idea that LPC species have acyl-chain-specific immunomodulatory effects, as well as the hypothesis that saturated LPC species are generally inflammatory, while unsaturated species are generally antiinflammatory^34^.

Neutrophils are increasingly recognized as critical regulators of tumor immunity, capable of both promoting and constraining tumor growth depending on their activation state and metabolic programming^37^. Tumor associated neutrophils can assume an anti-tumor N1 phenotype that produce cytotoxic ROS and support T-cell responses, or a pro-tumor N2 phenotype that promote angiogenesis, immunosuppression, and metastasis^38^. While broad neutrophil suppression (such as corticosteroid treatment) can alleviate ICB-irAEs, it carries the cost of dampening systemic anti-tumor immunity^39,40^. Specifically, there is an unmet need for studies monitoring intratumoral myeloid populations after systemic corticosteroids; lower NLR correlating with improved outcomes in ICB-treated patients on prolonged systemic corticosteroids supports a tumor associated neutrophil-dependent microenvironment in these patients^41^. A more precise approach would selectively induce intrinsic apoptosis in hyperactivated neutrophils, allowing physiological resolution without global immunosuppression.

Our data indicate that LPC 18:2 elicits a distinct neutrophil fate characterized by mitochondrial depolarization, cytochrome c release, and caspase-3/7 activation consistent with intrinsic apoptosis, occurring concurrently. We hypothesize that this combination serves as a two-phase pro-resolving program: transient ROS may aid microbial and tumoricidal functions, whereas subsequent ROS-independent apoptosis commits neutrophils to controlled clearance. Apoptotic neutrophils are efficiently efferocytosed by macrophages, promoting an anti-inflammatory phenotype via TGF-β, IL-10, and specialized pro-resolving mediators^42,43^. Given that colonic infiltration of neutrophils and epithelial damage are observed in ICB-induced colitis^44^, LPC 18:2-driven apoptotic remodeling could limit mucosal injury by facilitating macrophage-mediated resolution. Thus, ROS-concurrent apoptosis may represent a therapeutic equilibrium that restrains ICB-irAE-associated inflammation while preserving anti-tumor immune competence of ICB antibodies.

Mechanistically, our data show that LPC 18:2 induced ROS production and apoptosis require lipid rafts, cholesterol-rich membrane microdomains. Disruption of lipid rafts through cholesterol depletion completely abrogated LPC 18:2-induced ROS and apoptosis. The requirement of intact lipid rafts for LPC 18:2 signaling is consistent with previous studies demonstrating raft-dependent signaling by LPC 16:0 in other cell-based models^31,32,45,46^. To our knowledge, this is the first evidence of unsaturated LPC species signaling also being dependent on intact lipid rafts. These results suggest that in human neutrophils, LPC 18:2 either engages a raft-resident receptor or directly incorporates into a cell membrane to induce its effects. Additional work will be needed to define the precise molecular intermediates linking raft signaling to mitochondrial apoptosis.

We previously observed that both LPC 16:0 and 18:2 associated with protection from ICB-irAEs, as patients who maintained higher circulating levels of either lipid were less likely to develop severe toxicities^9^. This shared trend suggests that abundant circulating LPCs mark a more homeostatic, less inflammatory immune metabolic state. However, only LPC 18:2 showed a specific inverse relationship with peripheral neutrophil counts and ICB-irAE severity, indicating an active role in restraining neutrophil-driven pathology. Here, *in vitro*, LPC 18:2, unlike LPC 16:0, induced a transcriptionally silent, intrinsic apoptotic program in human neutrophils characterized by cytochrome c release and caspase-3/7 activation concurrent with ROS generation. In ICB-treated mice, LPC 18:2 supplementation limits neutrophilia and protects against colitis, whereas LPC 16:0 fails to do so^9^. This specificity likely reflects structural and signaling differences between LPC species. Unsaturated LPC 18:2 can promote pro-resolving macrophage efferocytosis, enhancing clearance of apoptotic neutrophils and release of anti-inflammatory mediators such as IL-10, TGF-β, and specialized pro-resolving lipid mediators^47^. In parallel, LPC 18:2 may sustain localized oxidative cues that reinforce caspase-3/7-dependent intrinsic apoptosis, ensuring activated neutrophils undergo controlled death rather than secondary necrosis. This coupled process of apoptosis followed by efficient efferocytic removal could create a feed-forward loop that restores mucosal integrity and terminates inflammation. Saturated LPC 16:0 lacks enzymatic versatility. Saturated LPC 16:0 is a poor substrate for further lipid modification, while unsaturated LPC 18:2 can be enzymatically remodeled or oxidized by lipoxygenases, cytochrome P450s, and acyltransferases to yield bioactive derivatives such as 13- and 9-hydroxyoctadecadienoic acids (HODEs) or re-enter phosphatidylcholine pools via LPCAT-mediated reacylation^48–51^. These metabolites participate in anti-inflammatory signaling, PPARδ activation, and membrane lipid remodeling, which are features that are absent in saturated species^52–56^. Thus, this provides LPC 18:2 greater biochemical “versatility” to engage pro-resolving and antioxidant pathways.

This study has several limitations. First, all experiments were performed *in vitro* using isolated primary human neutrophils, which may not fully capture the complex signaling environment present *in vivo*. In circulation, LPCs are largely bound to albumin^53^, which may influence their activity; this was not examined here. Additionally, while we examined the two most abundant LPC species in the plasma^17^, other LPC species may exert additional, context-dependent effects on neutrophils and other immune subsets. Finally, although our data support a cholesterol- and lipid raft–dependent mechanism, the downstream molecular mediators remain undefined. Addressing these gaps will be critical to establish the physiological and therapeutic relevance of LPC–neutrophil interactions.

In summary, our findings identify LPC 18:2 as a transcriptionally quiet lipid mediator that promotes neutrophil apoptosis through cholesterol- and lipid raft–dependent signaling, in contrast to LPC 16:0, which activates proinflammatory transcriptional programs and induces inflammatory cell death.

These findings highlight the functional diversity of lysophospholipids and underscore their potential as regulators of immune cell fate. More broadly, this work suggests that treatment with specific LPC species may represent a novel therapeutic approach for controlling pathological neutrophil activation in inflammatory conditions including ICB-irAEs.

## 5 Data Availability Statement

The bulk RNA-seq dataset generated for this study will be available starting March 6^th^, 2026 at Gene Expression Omnibus under accession number GSE311051 (http://www.ncbi.nlm.nih.gov/geo/).

## 6 Author Contributions

PS and SS conceptualized and designed the study. PS, AG and SS were responsible for methodologies, data interpretation and data curation. PS, AG, MC, NN and CF conducted neutrop il experiments. The original draft of the manuscript was written by PS and AG. MC performed RNA preparation for bulk sequencing and contributed to data visualization, figure organization, and graphical refinement. SS, MC, AG, IM, MV and MR contributed to reviewing and editing the manuscript. Funding acquisition and overall supervision of the project were provided by SS. All authors reviewed, edited, and approved of the final manuscript.

## 10 Acknowledgments

We acknowledge the use of the Sequencing Core Facility at the La Jolla Institute for Immunology (RRID:SCR_023107) for sequencing services. The NovaSeq 6000 used in this study was acquired through the NIH Shared Instrumentation Grant Program (S10), and we gratefully acknowledge its support (S10OD025052).

## 12 Figure Legends

**Supplemental Figure 1.**

(A) Gating strategy for human neutrophil cleaved caspase-3/7 assay.

(B) Gating strategy for human neutrophil ROS assay.

(C) Gating strategy for human neutrophil cytochrome c release assay.

(D) Gating strategy for human neutrophil MitoTracker assay.

