## Supplemental Figure 1 for "LPC 18:2-Driven Apoptosis In Neutrophils Is Non-Inflammatory and Lipid Raft Dependent"

### Supplementary Figure 1

#### A. Caspase Assay (Figure 1b, 3a, and 4c)

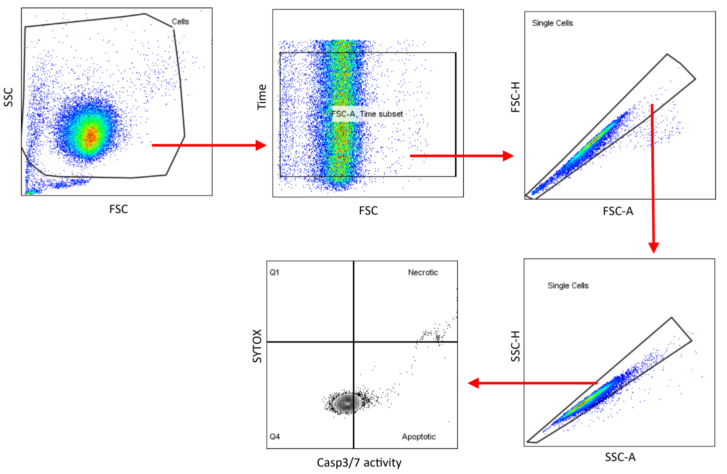


#### B. ROS Assay (Figure 1f and 4a)

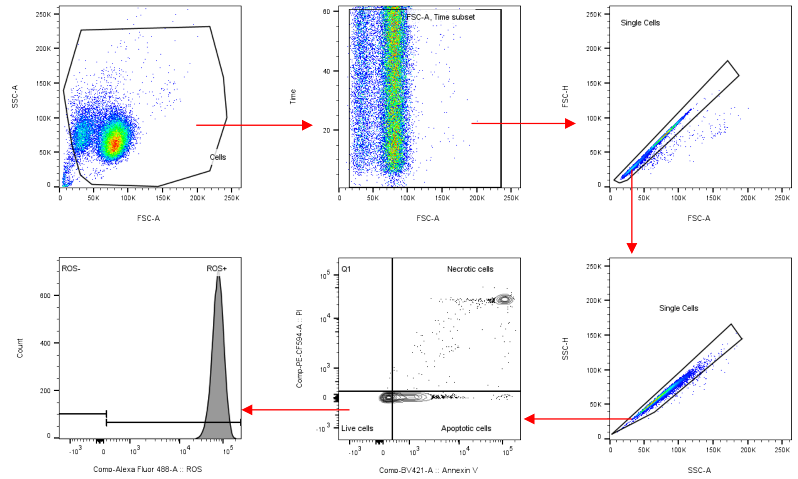


#### C. Cytochrome C (Figure 3b)

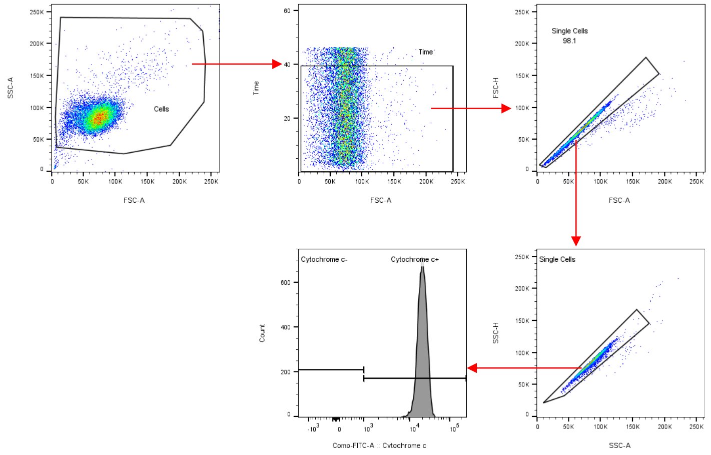


#### D. MitoTracker Assay (Figure 3c and 3d)

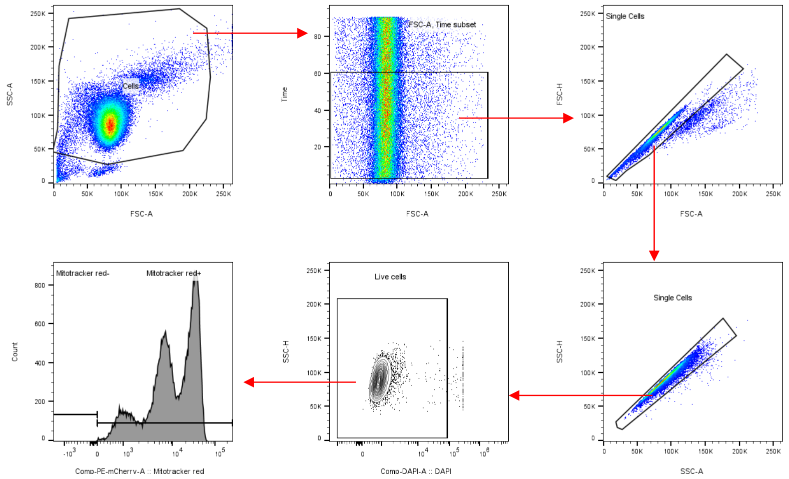
